# Effects of underlying gene-regulation network structure on prediction accuracy in high-dimensional regression

**DOI:** 10.1101/2020.09.11.293456

**Authors:** Yuichi Okinaga, Daisuke Kyogoku, Satoshi Kondo, Atsushi J. Nagano, Kei Hirose

**Affiliations:** Graduate School of Mathematics, Kyushu University, 744 Motooka, Fukuoka 819-0395, Japan; The Museum of Nature and Human Activities, 6 Yayoigaoka, Sanda, Hyogo 669-1546, Japan; Agriculture and Biotechnology Business Division, Toyota Motor Corporation, Miyoshi, Aichi 470-0201, Japan; Faculty of Agriculture, Ryukoku University, Otsu, Shiga 520-2194, Japan; Institute of Mathematics for Industry, Kyushu University, 744 Motooka, Fukuoka 819-0395, Japan and RIKEN Center for Advanced Intelligence Project, 1-4-1 Nihonbashi, Chuo-ku, Tokyo 103-0027, Japan

## Abstract

**Motivation:** The least absolute shrinkage and selection operator (lasso) and principal component regression (PCR) are popular methods of estimating traits from high-dimensional omics data, such as transcriptomes. The prediction accuracy of these estimation methods is highly dependent on the covariance structure, which is characterized by gene regulation networks. However, the manner in which the structure of a gene regulation network together with the sample size affects prediction accuracy has not yet been sufficiently investigated. In this study, Monte Carlo simulations are conducted to investigate the prediction accuracy for several network structures under various sample sizes.

**Results:** When the gene regulation network was random graph, the simulation indicated that models with high estimation accuracy could be achieved with small sample sizes. However, a real gene regulation network is likely to exhibit a scale-free structure. In such cases, the simulation indicated that a relatively large number of observations is required to accurately predict traits from a transcriptome.

**Availability and implementation:** Source code at https://github.com/keihirose/simrnet

## 1 Introduction

Technological advancements have enabled the collection of highly multidimensional data from biological systems (Gehlenborg *et al*., 2010; Mochida and Shinozaki, 2011; Li and Sillanpää, 2012; Hasin *et al*., 2017). For example, RNA sequencing quantifies expression levels of thousands of genes. Such omics data is useful in predicting organismal traits, with empirical applications including diagnosis and classification of diseases and prediction of patient survival (van’t Veer *et al*., 2002; Bøvelstad *et al*., 2007; Chan *et al*., 2016; Nandagopal *et al*., 2019) and possible future applications in predicting crop yields (Kremling *et al*., 2018), insecticide resistance (Dermauw *et al*., 2013), and environmental adaptation (Nagano *et al*., 2019).

A common challenge in predicting traits from omics data is the dimension of the data far exceeding that of the sample size (known as high-dimensional regression). For example, if one is to apply least-squares estimation in multiple regression (e.g. trait ≈ *β*_0_ + *β*_1_gene_1_ + *β*_2_gene_2_ + ⋯) to predict a trait value from a transcriptome, the sample size needs to be (at least) larger than the number of model parameters. However, because transcriptome studies typically observe thousands of genes, a sample size exceeding the number of genes is not realistic at present. In this case, high-dimensional regression modeling must be considered.

The least absolute shrinkage and selection operator (lasso, Tibshirani, 1996) is one of the most frequently used methods for high-dimensional regression. It simultaneously achieves variable selection and parameter estimation. Theoretically, the prediction accuracy of the lasso is highly dependent on the correlation structure among exploratory variables; it is high under certain strong conditions, such as the compatibility condition (van de Geer and Buhlmann, 2009). However, in practice, it is not easy to check whether the compatibility condition holds. Another popular estimation method for high-dimensional regression is principal component regression (PCR, Jolliffe, 1986). PCR is a two-stage procedure: first, principal component analysis is conducted for predictors, following which the regression model on which the principal components are used as predictors is fitted. This method may perform well when the exploratory variables are highly correlated.

It is reasonable to assume that gene regulation networks will result in conditional independence among the levels of gene expression (Wei and Li, 2007; Dobra *et al*., 2004; Yu *et al*., 2013). Here, two variables are conditionally independent when they are independent given other variables (e.g. two focal variables are independently influenced by a third variable, Wille and Buhlmann, 2006). When a random vector of exploratory variables follows a multivariate normal distribution, two variables are conditionally independent if and only if the corresponding element of the inverse covariance matrix is nonzero. Essentially, the networks are characterized by the nonzero pattern of the inverse covariance matrix.

One of the most notable characteristics of biological networks is their scale-free nature, that is, the degree distribution of the network follows a power-law expressed as *p*(*x*) ∝ *x*^−*γ*^ (*γ* > 1) (Barabási and Albert, 1999; Milo *et al*., 2002). Empirical studies suggest that biological networks are often scale-free (Barabasi and Oltvai, 2004; Albert, 2005; Arita, 2005), although exceptions have also been found Broido and Clauset (2019). Therefore, it is reasonable to consider the problem of high-dimensional regression when the networks of exploratory variables are scale-free. Here, it should be noted that the relative performance of different high-dimensional regression techniques may depend on sample sizes. However, to the best of our knowledge, the effect of the gene regulation network structure together with sample size on prediction accuracy has not yet been sufficiently investigated.

This paper provides a general simulation framework to study the effects of correlation structure in explanatory variables. As an example, the prediction of ambient temperature from the transcriptome, for which good empirical data is available (Nagano *et al*., 2012, 2019), is considered. It should be noted that the implementation of the proposed procedure is independent of the empirical data in Nagano *et al*. (2012, 2019); the proposed framework may be applied to predict any consequence of gene expression differences (e.g. crop yield). The proposed framework is based on the Monte Carlo simulations. Three datasets of transcriptome and their traits are generated. The datasets are characterized by the covariance structure of exploratory variables; one of the covariance structures corresponds to the scale-free gene regulation network. Both lasso and PCR are applied to these simulated datasets to investigate the prediction accuracy with different types of gene regulation networks. The sample size is also varied to examine its effect on the prediction accuracy.

The remainder of this paper is organized as follows. Section 2 describes prediction methods for high-dimensional regression in the given simulation. Section 3 discusses the proposed simulation framework. Finally, Section 4 presents the concluding remarks.

## 2 Prediction methods for high-dimensional data

Suppose that we have *n* observations {(***x**_i_, y_i_*) | *i* = 1,…, *n*}, where ***x**_i_* are *p*-dimensional vector of explanatory variables and *y_i_* are responses (*i* = 1,…, *n*). Let *X* = (***x***_1_,⋯, ***x***_*n*_)^*T*^ and *Y* = (*y*_1_,⋯, *y_n_*)^*T*^. Consider the linear regression model:

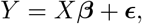

where ***ϵ*** = (*ϵ*_1_,⋯, *ϵ_n_*)^*T*^ is a vector of error variables with *E*(***ϵ***) = **0** and var(***ϵ***) = *σ*^2^*I_n_*.

### 2.1 Lasso

The lasso minimizes a loss function that consists of quadratic loss with a penalty based on an *L*_1_ norm of a parameter vector:

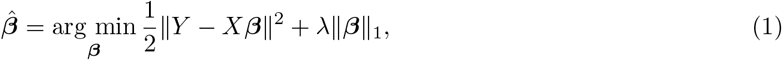

where λ > 0 is a regularization parameter. Because of the nature of the *L*_1_ norm in the penalty term, some of the elements of the coefficients are estimated to be exactly zero. Thus, variable selection is conducted, and only variables that correspond to nonzero coefficients affect the responses.

### 2.2 PCR

In some cases, the first few largest eigenvalues of the covariance matrix of predictors (i.e., proportional contributions of principle components) can be considerably large (e.g., spiked covariance model, Johnstone *et al*., 2001). In such a case, the lasso may not function effectively in terms of both prediction accuracy and consistency in model selection, because the conditions for its effective performance (e.g., compatibility condition, Bühlmann and van de Geer, 2011) may not be satisfied. This issue could be addressed using PCR because it transforms data with a large number of highly correlated variables into a few principal components. In the first stage of PCR, principal component analysis is applied to predictors, and the dimension of ***x***_*i*_ is reduced to *d* (*d* ≪ *p*). In this work, *d* was chosen such that *d* principle components collectively explain 90% or more variance (and *d* − 1 principle components do not). Then, in the second stage, regression analysis is conducted, for which the principal components are used as predictors. Here, the regression coefficients in the second stage are estimated by the lasso.

## 3 Simulation framework

An overview of the simulation is presented in Fig. 1. First, the model that defines the relationship between the trait and the levels of gene expression was parameterized. This was done using the empirical data in Nagano *et al*. (2019), which quantified the transcriptome of wild *Arabidopsis halleri* subsp. *gemmifera* weekly for two years in their natural habitat as well as bihourly on the equinoxes and solstices (*p* = 17205 genes for *n* = 835 observations). Three types of simulated data were generated using different covariance matrices of genes, denoted as Σ_*j*_ (*j* = 1, 2, 3). Σ_1_ is the sample covariance matrix of genes. Generally, none of the elements of the inverse of sample covariance matrix are exactly zero, implying that each gene interacts with all the other genes. Such a fully connected network is ineffective in terms of interpretation of the mechanism of gene regulation. Thus, two other covariance matrices were produced to simulate sparse networks based on the sample covariance matrix Σ_1_. Σ_2_ is generated by the graphical lasso (Yuan and Lin, 2007), which corresponds to the random graph. Although the graphical lasso is widely used because of its computational efficiency, real networks are often scale-free. Therefore, Σ_3_, which corresponds to the scale-free network, was generated here. The estimation of scale-free networks is achieved by the reweighted graphical lasso (Liu and Ihler, 2011). Based on these three covariance matrices Σ_*j*_ (*j* = 1, 2, 3), the simulated transcriptome data were generated from the multivariate normal distribution. The simulated ambient temperature were generated from simulated transcriptome data. Finally, lasso and PCR were applied to these simulated data to compare their prediction accuracies. The sample sizes of the simulated data were varied to investigate the relationship between prediction accuracy and sample sizes.

**Figure 1:**
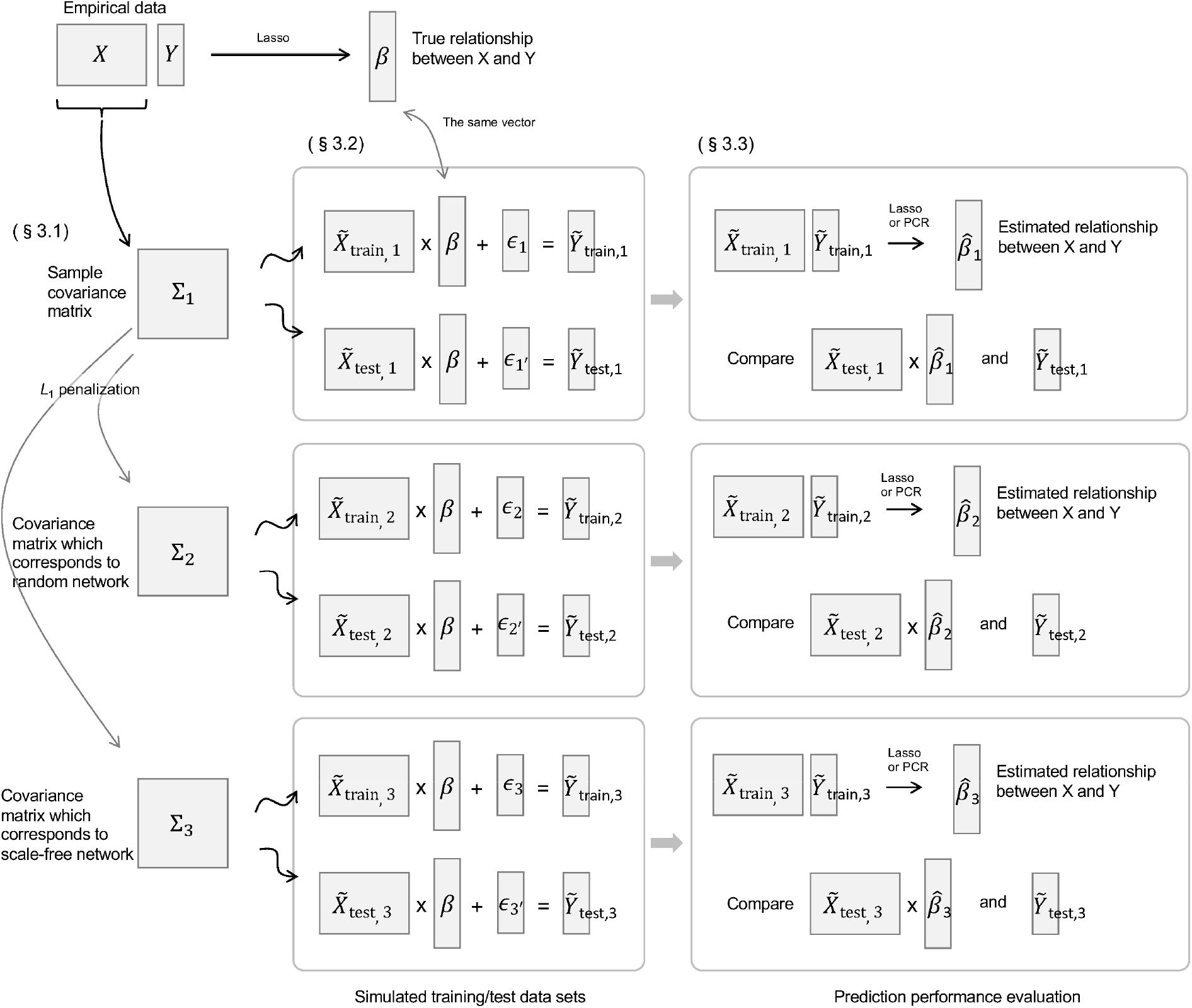
Overview of the simulation.

### 3.1 Evaluation of the estimation procedure

The performance of the estimation procedure is investigated by the following expected prediction error:

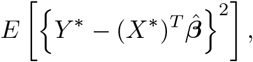

where *X*\* and *Y*\* follow *X*\* ~ *N*(**0**, Σ_*j*_) (*j* = 1, 2, or 3) and *Y*\* ~ *N*((*X*\*)^*T*^ ***β***, *σ*^2^), respectively. The estimator 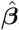 is obtained using current observations, while *X*\* and *Y*\* correspond to future observations. The Σ_*j*_ (*j* = 1, 2, 3), ***β***, and *σ*^2^ are true values but unknown. In practice, these parameters are defined by using the actual dataset, (*X, Y*). Detail of setting of these parameters will be presented in Section 3.2.

To estimate the expected prediction error, the Monte Carlo simulation is conducted. We first randomly generate training and test data, 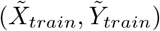 and 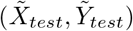, respectively. Here, 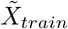 follows a multivariate normal distribution with mean vector *μ_X_* and variance-covariance matrix Σ_*j*_, where *μ_X_* is the sample mean of *X*. Then, 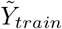 is generated by 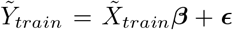, where ***ϵ*** is a random sample from *N*(**0**, *σ*^2^*I*) with *I* being an identity matrix. The test data, 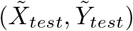, are generated by the same procedure as 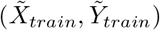 but independent of 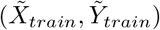. The number of observations for the training and test data are *N* (*N* = 50, 100, 200, 300, 500, 1000) and 1000, respectively. The lasso and the PCR described in Section 2 are performed with training data 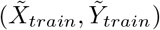, following which RMSE is calculated in (8). The above process, from random generation of data to RMSE calculation, was performed 100 times.

### 3.2 Parameter setting

#### 3.2.1 Covariance structures

Here, the characterization of the network structure of predictors by conditional independence is considered. When the predictors follow a multivariate normal distribution, the network structure based on the conditional independence corresponds to the nonzero pattern of the inverse covariance (precision) matrix. In other words, the network structure is characterized by the inverse covariance matrix of predictors.

Let *S* be the sample covariance matrix of predictors, that is, 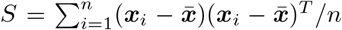 with 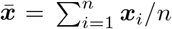. Let 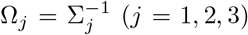. Σ_1_ is a ridge estimator of the sample variance-covariance matrix, that is, Σ_1_ = *S* + *δI*. Here *δ* is a small positive value (in this simulation, *δ* = 10^−5^). The term *δI* allows the existence of Ω_1_. Note that because Ω_1_ is not sparse, it leads to the complete graph, which is of no use in interpreting gene regulatory networks. To generate a covariance matrix whose inverse matrix is sparse, *L*_1_ penalization is employed for the estimation of Ω_2_ and Ω_3_ as follows:

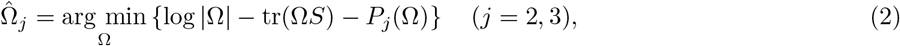

where *P_j_*(Ω) (*j* = 2, 3) are penalty terms which enhance the sparsity of the inverse covariance matrix. To estimate the sparse inverse covariance matrix, the lasso penalty is typically used as follows:

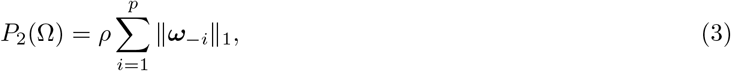

where 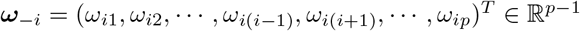. The problem (3) is referred to as the graphical lasso (Yuan and Lin, 2007), and there exists several efficient algorithms to obtain the solution (Friedman *et al*., 2008; Witten *et al*., 2011; Boyd, 2011). The estimator of (2) with (3) corresponds to Ω_2_ and 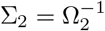.

The lasso penalty (3) does not enhance scale-free networks. It penalizes all edges equally so that the estimated graph is likely to be a random graph, that is, the degree distribution becomes a binomial distribution. To enhance scale-free networks (i.e., power-law distribution), the log penalty (Liu and Ihler, 2011) is used as follows:

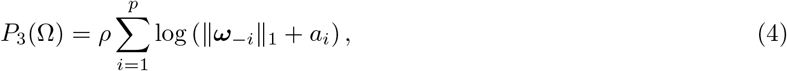

where *a_i_* > 0 are tuning parameters. From a Bayesian viewpoint, the prior distribution which corresponds to the log penalty becomes the power-law distribution (Liu and Ihler, 2011); thus, the penalty (4) is likely to estimate the scale-free networks. The estimator of (2) with (4) corresponds to Ω_3_.

Because the log-penalty (4) is nonconvex, it is not easy to directly optimize (2). To implement the maximization problem (2), Liu and Ihler (2011) constructed the minorize-maximization (MM) algorithm (Hunter and Lange, 2004), in which the weighted lasso penalty 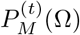 with current parameter 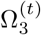 is used:

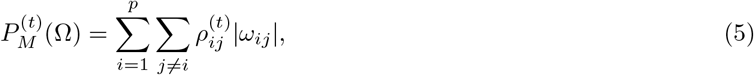

where 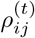 are the weights

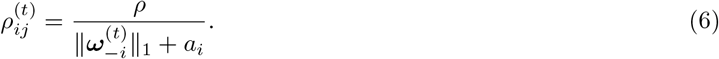

Because the weighted graphical lasso can be implemented by a standard graphical lasso algorithm, the estimator is obtained as the following algorithm:

1. Set *t* = 0. Get 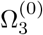 via ordinary graphical lasso. Repeat 2 to 4 until convergence.
2. Update weights 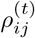 using (6).
3. Get 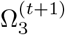 via the weighted graphical lasso (2) with penalty (5).
4. *t* ← *t* + 1.

To obtain Σ_2_ and Σ_3_, the tuning parameters *a_i_* (*i* = 1…, *p*) and *ρ* must be determined. Following the experiments in Liu and Ihler (2011), *a_i_* = 1 was set for *i* = 1…, *p*. To select the value of the regularization parameter *ρ*, several candidates were first prepared. In our simulation, the candidates were *ρ* = 0.3, 0.4,0.5, 0.6,0.7. From these, the value of *ρ* was selected such that the extended Bayesian information criterion (EBIC, Chen and Chen, 2008; Foygel and Drton, 2010)

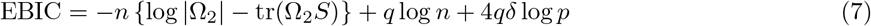

was minimized. Here, *q* is the number of nonzero parameters of the upper triangular matrix of 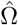, and *δ* ∈ [0,1) is a tuning parameter. As the value of *δ* increases, a sparser graph is generated. Note that *δ* = 0 corresponds to the ordinary BIC (Schwarz, 1978). We set *δ* = 0.5 because Foygel and Drton (2010) showed that *δ* = 0.5 yielded good performance in both simulated and real data analyses. As a result, the EBIC selected *ρ* = 0.5.

The upper triangular matrix Ω_3_ must be estimated with the reweighted graphical lasso problem. A value of *p* = 17205 results in *p*(*p* + 1)/2 ≈ 148 million parameters. As a result, with the machine used in this study (Intel Core Xeon 3 GHz, 128 GB memory), it would take several days to conduct the reweighted graphical lasso approach, even with a small number of iterations such as *T* = 5. For this reason, *T* = 5 iterations were employed to produce Σ_3_ here.

Fig. 2 depicts the logarithm of the largest 30 eigenvalues of Σ_*j*_ (*j* = 1, 2, 3). The first few largest eigenvalues of Σ_3_ are significantly larger than those of Σ_2_, implying that the scale-free networks tend to produce predictors with large correlations.

**Figure 2:**
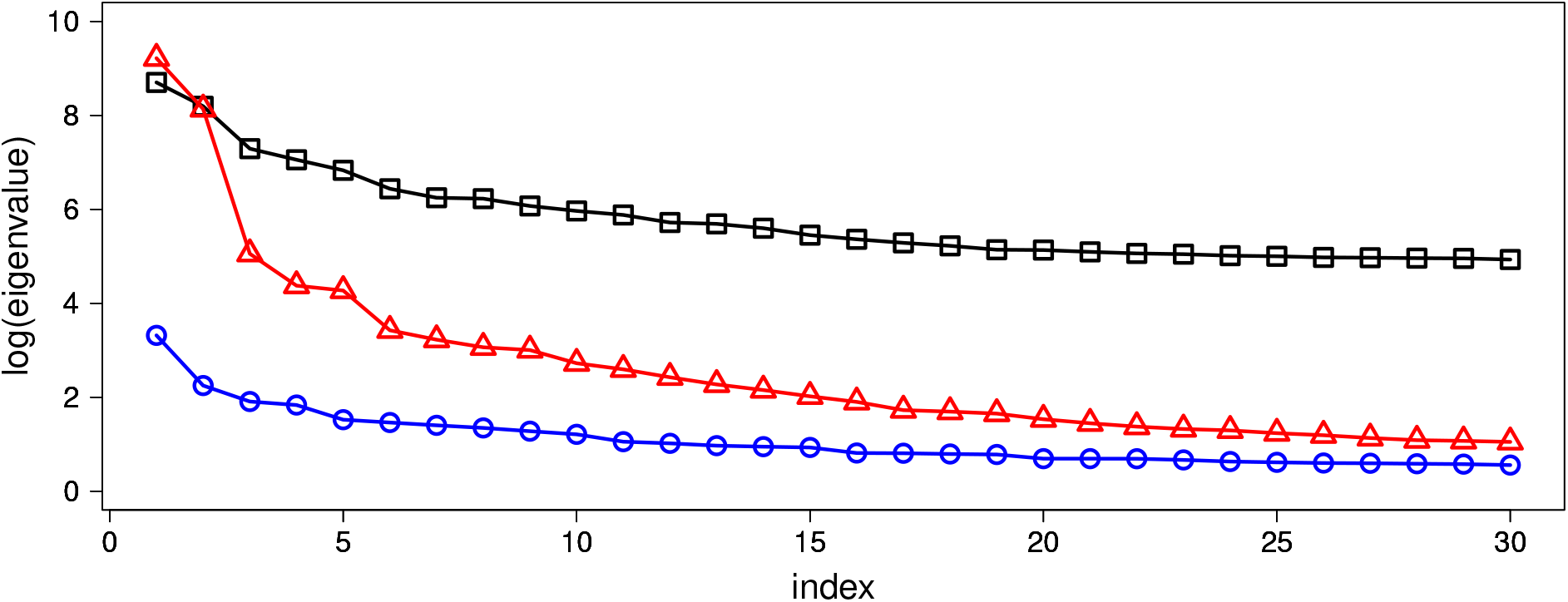
Logarithm graph of the largest 30 eigenvalues of Σ_1_ (black square), Σ_2_ (blue triangle) and Σ_3_ (red circle). The horizontal axis expresses the index of eigenvalues arranged in descending order.

#### 3.2.2 Regression parameters

The values of ***β*** and *σ*^2^ are determined as follows. First, 10-fold cross-validation is performed as described below, and the regularization parameter λ in (1) is selected. The data (*X, Y*) are divided into ten datasets, (*X*^(*j*)^, *Y*^(*j*)^) (*j* = 1,…, 10), which consist of almost equal sample sizes. Let *X*^(−*j*)^ = (*X*^(1)^,…, *X*^(*j*−1)^, *X*^(*j*+1)^,…, *X*^(10)^), and *Y*^(−*j*)^ = (*Y*^(1)^,…, *Y*^(*j*−1)^, *Y*^(*j*+1)^,…, *Y*^(10)^) (*j* = 1,…, 10). For each *j* (*j* = 1,…, 10), the training and test data are defined by (*X*^(−*j*)^, *Y*^(−*j*)^) and (*X*^(*j*)^, *Y*^(*j*)^), respectively. Then, the parameter 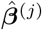 (*j* = 1,…, 10) is found by the lasso:

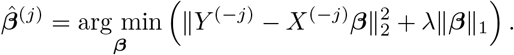

For each *j*(*j* = 1,…, 10), the verification error is calculated as follows:

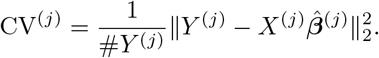

Then, λ is adopted such that it minimizes 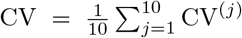, the mean of CV^(*j*)^. Following this, the dataset (*X, Y*) is again randomly divided into two datasets: test data (*X_test_, Y_test_*) and training data (*X_train_, Y_train_*). Lasso estimation (1) is performed using the training data, with λ obtained by the above 10-fold cross-validation. Then, ***β*** is defined as the lasso estimator, resulting in the number of nonzero parameters of ***β*** being 259. Fig. 3 shows the histogram of nonzero parameters of ***β***. It is seen that the majority of the nonzero coefficients were close to zero; only 15 parameters had absolute values larger than 0.1.

**Figure 3:**
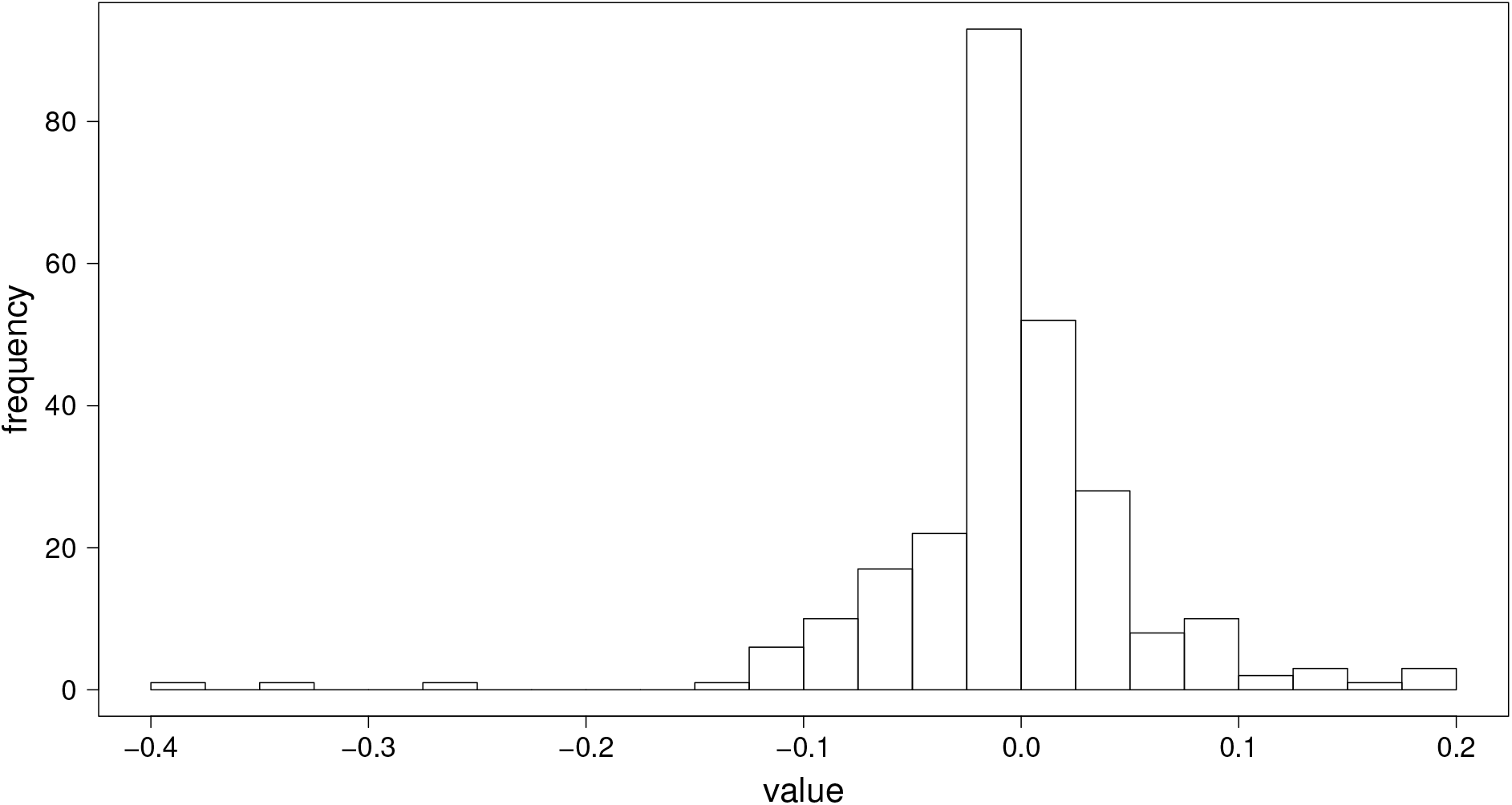
Histogram of 259 nonzero parameters of ***β***.

In addition, the root mean squared error (RMSE) is calculated as follows:

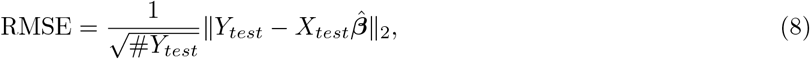

and the variance of errors, *σ*^2^, is defined by *σ*^2^ = (RMSE)^2^.

### 3.3 Results

The box and whisker plot of the RMSE is drawn in Fig. 4. The horizontal axis is *N* (the number of observations of training data) and the vertical axis is the RMSE based on 1000 observations of test data.

**Figure 4:**
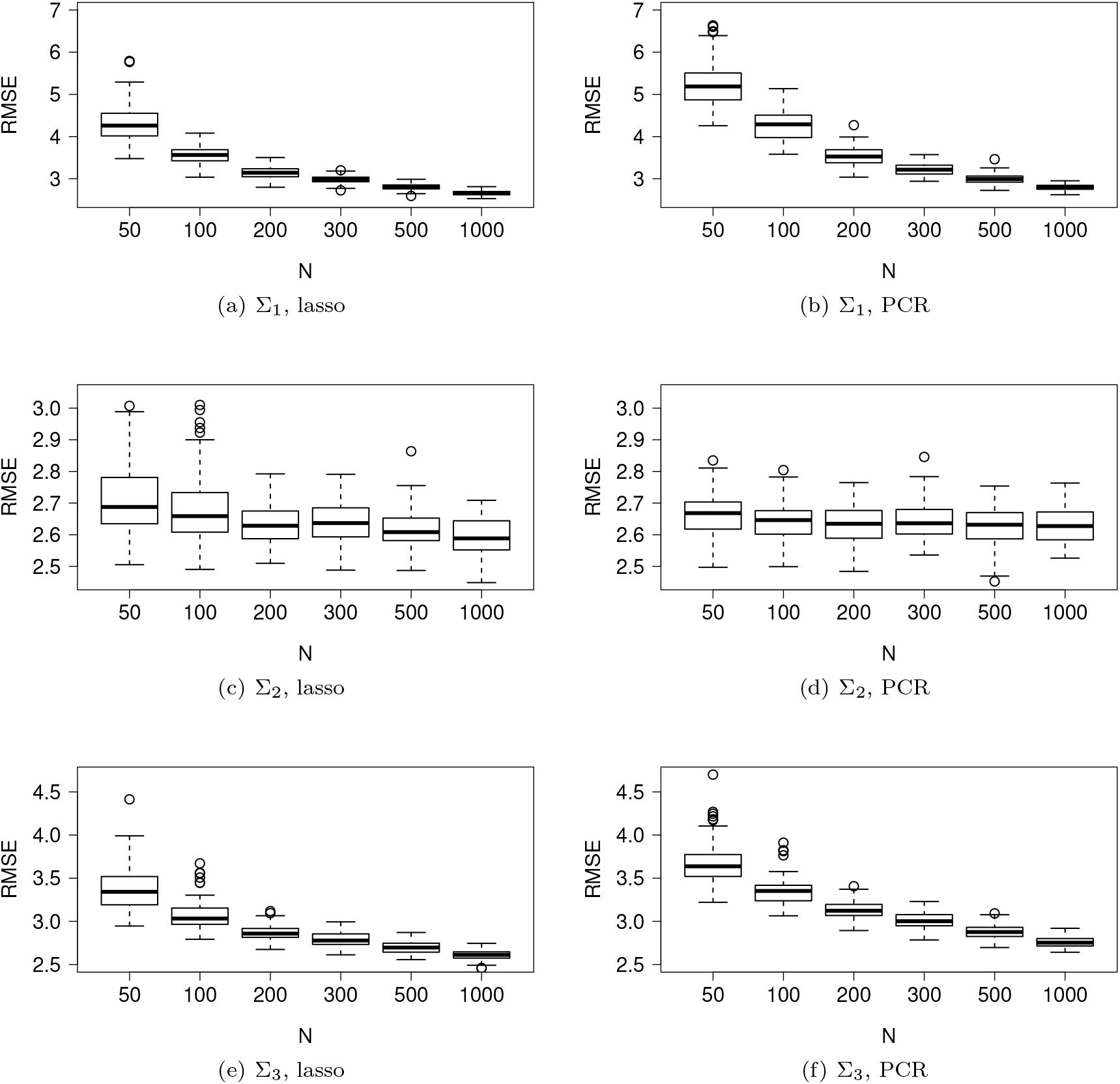
Box and whisker plot of RMSE. The variance-covariance matrix used in the simulations is Σ_1_ in (a)–(b), Σ_2_ in (c)–(d), and Σ_3_ in (e)–(f). The regression model is estimated by the lasso (Section 2.1) in (a), (c), and (e) and by PCR (Section 2.2) in (b), (d), and (f).

We compared the performance of the lasso with that of the PCR. When Σ_1_ and Σ_3_ were used, the PCR performed worse than the lasso for small sample sizes. Some predictors associated with small eigenvalues may affect prediction performance. Meanwhile, for Σ_2_, the performance of PCR was slightly more stable than that of the lasso for small sample sizes.

The prediction accuracy was compared among the three covariance structures. For both lasso and PCR, when Σ_1_ and Σ_3_ were used, the values of RMSE decreased as *N* increased. On the other hand, when Σ_2_ was used, the values of RMSE remained almost unchanged, approximately ranging between 2.5–3, as *N* increased. In the case of scale-free (Σ_3_), when the number of observations was large, the estimation accuracy was approximately between 2.5–3, which was almost identical to the accuracy of Σ_2_. For example, the mean RMSE at *N* = 1000 in Fig. 4 (e) was nearly equal to that at *N* = 50 in (c). The reason the accuracy of Σ_2_ remained high even at *N* = 50 is considered to be that Σ_2_ was weakly-correlated (Fig. 2) and the majority of the nonzero parameters of ***β*** were small (Fig. 3). Such a weakly-correlated covariance matrix implies conditions on Σ_2_ that achieve nearly optimal rates may be satisfied (e.g., van de Geer and Buhlmann, 2009).

As described before, Σ_1_ was the sample covariance matrix, while Σ_3_ (and Σ_2_) was estimated using the graphical lasso. As the lasso-type regularization methods shrink parameters toward zero, the correlations among exploratory variables reduce with the graphical lasso. Therefore, Σ_3_ resulted in smaller correlations as compared to Σ_1_. Consequently, the prediction accuracy reduces with stronger correlations.

### 3.4 Code availability

The proposed simulation is implemented in `R` package `simrnet`, which is available at https://github.com/keihirose/simrnet. Below is a sample code of the `simrnet` in `R`:

~~~
library(devtools)
install_github(“keihirose/simrnet”) #install package
library(simrnet) #load package
data(nagano2019)
attach(nagano2019)
rho <- (1:9) / 10 #tuning parameters for glasso
pars <- genpar(X,Y,rho) #set true parameter
result <- simrnet(pars, times.sim=100) #conduct simulation
plot(result)
~~~

When *p* = 100, it took less than 12 minutes to conduct the simulation with 100 replications using the machine employed herein (Intel Core Xeon 3GHz, 128GB memory). For high-dimensional data such as *p* = 17205, which was used in the simulation presented in this paper, several days were required to complete the simulation task.

## 4 Concluding remarks

In a gene regulation network, a gene regulates a small portion of a genome, not all the genes in a genome. This indicates that gene regulation network is expected to be a sparse network rather than a complete graph. Therefore, two covariance matrices indicating sparse networks (Σ_2_, Σ_3_) were prepared in addition to a covariance matrix derived from empirical data (Σ_1_). Generally, although hundreds of genes contribute to defining a trait, their contributions are not equal. It is frequently observed that genes regulating a trait include a few large-effect genes and several small-effect genes. This property was reflected in the distribution of ***β*** (Fig. 3). When a limited number of regression coefficients had a large contribution to the definition of a trait, and the gene regulation network was random (Σ_2_), the simulation indicated that models with high estimation accuracy could be developed from a small number of observations (Fig. 4). However, a real gene regulation network is likely to exhibit scale-free structure. In such cases, the simulation indicated that the prediction of traits from a transcriptome requires a relatively large number of observations to produce good performance (Σ_1_, Σ_3_, Fig. 4). In conclusion, it is necessary to secure sufficiently large sample sizes when performing regression analysis of data with scale-free network.

Conventional theory on the relationship between RMSE and sample size has been developed under the assumption that the sample size exceeds the number of exploratory variables (e.g., Fahrmeir *et al*., 2007). However, omics data, which is rapidly being accumulated, results in high dimensional data with strong correlations. Thus, our simulation study considered more complicated settings than the traditional ones. Our simulation, or its extension, may be used in the future to find clues about theoretical aspects that may ultimately lead to the development of a sample size determination technique for omics data. Another important future research topic is the development of methods that have better estimation accuracy than the lasso in the case of small sample sizes.

## Acknowledgements

The authors would like to thank Mr. Kanta Miura for the valuable discussions.

